# SALL4 is required for *YAP1*-dependent malignant and regenerative hepatocyte reprogramming into cholangiocyte lineage

**DOI:** 10.1101/2025.08.15.670568

**Authors:** Minwook Kim, Yoojeong Park, Rachel Covitz, Joseph Kwon, Jia-Jun Liu, Silvia Liu, Sungjin Ko

## Abstract

Hepatocytes (HCs), which share a developmental origin with cholangiocytes (CCs), have the capacity to undergo reparative reprogramming into CCs in response to liver injury and, under specific conditions, can also transform malignantly into cholangiocarcinoma (CCA). However, the molecular mechanisms governing HC plasticity in liver diseases remain poorly understood. In this study, we investigated the role of *Spalt Like Transcription Factor 4 (SALL4)*, an oncofetal transcription factor, in both malignant and regenerative HC fate transitions toward the biliary lineage. Using Sleeping Beauty hydrodynamic tail vein injection-mediated murine liver cancer models, we explored HC-to-CCA transformation, while the DDC diet-induced cholestasis model was used to investigate regenerative HC-to-CC reprogramming. Our findings reveal that SALL4 is specifically required for *myristoylated Akt (myrAkt)-YAP1*^*S127A*^ *(AY)*-driven HC-to-CCA transformation, as its loss significantly suppressed malignant reprogramming and clonal expansion. Surprisingly, SALL4 overexpression also prevented *AY*-driven CCA development while promoting the expansion of liver progenitor cell (LPC)-like fatty HCs. Mechanistically, we propose *Bmi1* as a key downstream effector of SALL4 in YAP1-dependent HC-to-CCA transformation. Additionally, in the DDC-fed cholestasis model, *Sall4* deletion enhanced HC-to-LPC activation while impairing LPC differentiation into mature CCs. These findings establish SALL4 as a critical regulator of HC plasticity in both malignant and regenerative contexts and highlight its potential as a therapeutic target for specific liver cancer subtypes.

**SIGNIFICANCE:** Hepatocyte plasticity supports repair but can drive malignancy, acting as a double-edged sword. We identify SALL4 as regulator of YAP1-driven hepatocyte-to-cholangiocyte reprogramming, revealing the YAP1–SALL4– BMI1 axis as a therapeutic target for cholangiocarcinoma.

## INTRODUCTION

Liver cancer is one of the most genetically and pathologically heterogeneous cancer types, with a devastating prognosis and a five-year survival rate of around 20%. It is traditionally classified into hepatocellular carcinoma (HCC) and cholangiocarcinoma (CCA), which account for approximately 80% and 20% of cases, respectively(1). Additionally, rare but intriguing liver cancer types, such as combined (cHCC-CCA) or mixed (mHCC/CCA) HCC-CCA, exhibit both hepatocellular and cholangiocellular differentiation within the same tumor or contain cells with intermediate morphology and/or expressing markers of both HCC and CCA, and share characteristics resembling liver progenitor cells (LPCs)(2-5).

The two hepatic main epithelial cell types, hepatocytes (HCs) and cholangiocytes (CCs) both originated from a common progenitor(6, 7), the hepatoblast, exhibit high degree of plasticity, allowing for interconversion between lineages under various liver diseases as needed(8, 9). In cholestatic diseases, reparative HC-to-CC conversion plays critical roles in regeneration by restoring damaged bile ducts when native CCs have lost their self-replication capacity. Notably, both murine and human studies have shown that HCs can undergo malignant reprogramming into CCA, a type of biliary cancer(10-12), highlighting the importance of understanding the liver cancer types in the context of their cellular origins. However, the molecular mechanisms that governs both reparative and malignant reprogramming of HCs toward the biliary lineage, despite sharing directional cues but leading to opposite outcomes, remain largely unknown.

Spalt Like Transcription Factor 4 (SALL4) is an oncofetal transcription factor essential for embryonic development and the maintenance of pluripotency in various progenitor and stem cells(13-16), including human embryonic stem cells (ESCs) and induced pluripotent stem cells (iPSCs). SALL4 plays a key role in fetal liver development(17), particularly in hepatic commitment and specification leading to hepatoblast formation. While its expression is nearly absent in the adult liver under normal conditions, recent studies suggest that SALL4 may have an unexplored role in liver cancer progression, particularly in specific aggressive HCC subtypes and hybrid HCC-CCA cases(18, 19).

Interestingly, SALL4 expression is elevated in poorly differentiated(3, 20, 21), KRT19^+^/EpCAM^+^ aggressive HCCs(19, 22), as well as in combined and mixed HCC-CCA tumors, but is only weakly detected in hepatoblastomas(23). This expression pattern in liver cancer with LPC characteristics suggests that SALL4 may be involved in hepatic dedifferentiation and plasticity, potentially influencing tumor development at specific disease stages. Moreover, conflicting reports on SALL4 expression in CCA(3, 20, 21, 24, 25) raise the possibility that its role has been underappreciated due to diagnostic challenges and tumor heterogeneity.

In this study, using established Sleeping Beauty hydrodynamic tail vein injection (SB-HDTVI)-mediated murine liver cancer models, we demonstrate that SALL4 is selectively required for *myristoylated Akt (myrAkt)-YAP1*^*S127A*^ *(AY)*-driven HC-to-CCA transformation and regenerative HC-to-CC reprogramming in the 3,5-diethoxycarbonyl-1,4-dihydrocollidine (DDC)-diet cholestasis model. We further show that SALL4 overexpression promotes the expansion of *AY*-transformed fatty HCs while preventing their malignant transition into CCA. Mechanistically, we suggest *Bmi1* as a functional downstream effector of SALL4 in YAP1-dependent HC-to-CCA transformation, phenocopying both gain- and loss-of-function effects of *Sall4*. These findings highlight SALL4 as a key regulator of hepatic plasticity and provide insights into its potential implications in liver cancer pathogenesis and therapeutic targeting.

## MATERIALS AND METHODS

### Animal

All animal care and experiments were conducted in accordance with the guideline of Institutional Animal Care and Use Committee (IACUC) at the University of Pittsburgh. *Sall4*^*flox/flox*^ mice were kindly provided by Dr. Masakazu Owashi (Kumamoto University, Japan) and *OPN-Cre*^*ERT2*^ mice generously shared by Dr. *Frédéric* Lemaigre (Université Catholique de Louvain, Belgium). To generate *Sall4*^*flox/flox*^;*OPN-Cre*^*ERT2*^ mice, *Sall4*^*flox/flox*^ mice were crossed with *OPN-Cre*^*ERT2*^ mice. Male mice were used for the *AY*-CCA model, while female mice were utilized for the *ANRAS*-mHCC/CCA and *KP53*-cHCC-CCA models. For DDC-diet, *Sall4* overexpression, and *Bmi1* deletion/overexpression experiments, FVB/NJ mice (Jax, 001800) were purchased and used.

### Hydrodynamic tail vein injection (HDTVI)

To induce tumorigenesis in the livers, we delivered *pT3-myrAkt, pT3-NRAS*^*G12V*^, *pT3-KRAS*^*G12D*^, *pT3-shTP53*, or *pT3-YAP1*^*S127A*^ plasmids via HDTVI(26), as previously described(10). All plasmids were extracted from 150-200 mL LB cultures of *E. coli* TOP10 (Invitrogen, C404003) using the NucleoBond^®^ Xtra Midi EF (TaKaRa, 740420.1) or ZymoPURE II Plasmid Midiprep Kit (Zymo Research, D4201), following the manufacturer’s instructions. Details on the plasmid quantities used for each tumor model are provided in **Supplementary Table 1**.

### DDC (3,5-Diethoxycarbonyl-1,4-dihydro-2,4,6-collidine) diet

To induce cholestatic injury in mice, we administered DDC diet (Inotiv, TD.230519). The diet contained 0.1% (wt/wt) of 3,5-Diethoxycarbonyl-1,4-dihydro-2,4,6-collidine (DDC, TCI America, D055825G) and was formulated using Teklad Rodent Diet 2020.

### AAV8 virus administration

Mice were injected intravenously with 1-2 × 10^12^ genome copies (GCs) of *adeno-associated virus serotype 8 (AAV8)* encoding *GFP* or *Cre* recombinase under the hepatocyte-specific thyroid-binding globulin (TBG) promoter. The following viral constructs were used: *AAV8-TBG-GFP* (Addgene, 105535-AAV8) and *AAV8-TBG-Cre* (Addgene, 107787-AAV8). Viruses were administered in 100 µL saline.

### Immunohistochemistry (IHC) and Immunofluorescence (IF)

Mouse livers were fixed in 10% buffered formalin (Fisher Scientific, 23-245685) for 48 hours and embedded in paraffin. Formalin-fixed, paraffin-embedded (FFPE) tissue samples were sectioned at 4 µm thickness using a microtome (Leica, RM2235) and mounted on positively charged slides (Azer Scientific, EMS200W+). Paraffin-embedded liver sections were deparaffinized using xylene (ThermoFisher, X-3S4) and rehydrated through sequential incubations in 100% and 95% ethanol (v/v) for 5 minutes each (repeated three time), followed by immersion in dH_2_O and washing in PBS. For antigen retrieval, heat-induced epitope retrieval (HIER) was performed using a pressure cooker for 20 minutes for HA-tag and □D-SMA, a microwave at 60% power for 12 minutes for panCK, V5-tag and SOX9, an incubation with Tris-EDTA buffer at 98°C for 20 minutes for CD45 or an autoclave with citrate buffer (pH 6.0) for HNF4A and Ki67. After antigen retrieval, the sections were allowed to cool to room temperature for 30 minutes and subsequently incubated in 3% H_2_O_2_ for 10 minutes to quench endogenous peroxidase activity. Following PBS washing, non-specific binding was blocked by incubating the sections in Super Block (ScyTek, AAA500) for 10 minutes at room temperature. Sections were then incubated overnight at 4°C with the primary antibody in a humidified chamber. After washing with PBS, sections were incubated with a biotinylated secondary antibody for 15 minutes at room temperature, followed by treatment with an avidin-biotin complex (Vector Laboratories, PK-6100) for 15 minutes, according to the manufacturer’s instructions. Diaminobenzidine (DAB) (Vector Laboratories, SK-4100) was used as a chromogen, and sections were counterstained with hematoxylin. The stained sections were dehydrated through an ascending ethanol series, cleared in xylene, and mounted with Cytoseal XYL (ThermoFisher, 8312-4). Primary and secondary antibodies used for IHC are listed in **Supplementary Table 2**.

For immunofluorescence, HIER was performed in a pressure cooker with Tris-EDTA (pH9.0) buffer for 20 minutes. Sections were subsequently washed in PBS, permeabilized for 5 minutes with PBS containing 0.25% Triton X-100 (Fisher Scientific Chemicals, A16046.AP) and blocked with PBS with 0.25% Triton X-100 and 10% bovine serum albumin (BSA) (Fisher BioReagents, BP9703100) for 45 minutes at room temperature. Following blocking, sections were incubated overnight at 4°C with primary antibodies diluted in PBS containing

0.25% Triton X-100 and 10% BSA. After incubation, sections were washed three times with PBS and incubated with fluorochrome-conjugated secondary antibodies in PBS 0.25% Triton X-100 and 10% BSA for 1h at room temperature. Sections were subsequently washed three times in PBS with 0.1% Triton X-100. For nucleus staining, Hoechst 33258 (Sigma-Aldrich, B-2883) was dissolved in dH_2_O at 1 mg/100 mL and applied to the sections for 1 minute, followed by rinsing with dH_2_O. Slides were mounted using Gelvatol mounting medium provided by the CBI at University of Pittsburgh. Confocal images were acquired using Nikon A1 confocal system. The primary and secondary antibodies used for IF are listed in **Supplementary Table 2**.

### Sirius Red staining

Picro-Sirius Red staining was performed using the Picro-Sirius Red Stain Kit (StatLab, KTPSRPT) according to the manufacturer’s protocol with slight modifications. Paraffin-embedded liver tissue sections were deparaffinized and rehydrated as described above. After rinsing in running tap water, the slides were immersed in Picro-Sirius Red solution for 60 minutes, rinsed briefly in 0.5% acetic acid 5 seconds each (repeated two times), and dehydrated in absolute ethanol for 10 seconds (repeated three times). Finally, sections were cleared in xylene for 1 minute each (repeated three times) and mounted with Cytoseal XYL (ThermoFisher, 8312-4).

### TUNEL Staining

Apoptotic cell death was detected using the ApopTag Plus Peroxidase *In Situ* Apoptosis Kit (Millipore, S7101) according to the manufacturer’s instructions. Briefly, deparaffinized and rehydrated liver tissue sections were treated with proteinase K, followed by incubation with the terminal deoxynucleotidyl transferase (TdT) enzyme to label DNA strand breaks. Signal development was performed using peroxidase-conjugated anti-digoxigenin antibody and DAB substrate. Slides were dehydrated and mounted for imaging.

### Hematoxylin and Eosin (H&E) staining

The tissue samples were processed overnight on a Lecia Peloris II tissue processor with a 12-hour processing schedule. The schedule consisted of one station of 70% ethanol (EtOH) (prepared by diluted 100% EtOH), two stations of 95% EtOH (Decon Labs, 801), three stations of 100% EtOH (Decon Labs, 2701), three stations of xylene (Leica, 3803665) and three stations of paraffin (Leica, 39602004). For reagents with multiple stations, the concentration was gradually increased, with the final station in each containing 100% pure reagent. Following processing, the samples were embedded in paraffin (Leica, Paraplast Plus). Tissue blocks were sectioned at 4 µm using a Leica RM2235 microtome. Sections were floated on a 44°C water bath and mounted onto Superfrost Plus slides (Thermo Fisher Scientific, 12-550-15). The slides were subsequently baked at 60°C for 30 minutes before staining. After baking, the slides were loaded onto a Leica ST5020 automate stainer with an integrated CV5030 coverslipper. H&E staining was performed using the Leica SelecTech staining system, which included Hematoxylin 560MX (3801575), Blue Buffer (3802918), Aqua Define MX (3803598) and Eosin 515 Phloxine (3801606).

### Image analysis

IHC brightfield images were acquired using the Evident Slideview VS200 (Olympus) with a 20x UPlanXApo objective (0.8 NA) and processed with Evident VS200 ASW 3.4.1 software. IHC images were analyzed and quantified using QuPath(27) and IF confocal images were analyzed and quantified using Fiji(28).

### Construct overexpression vectors

For the overexpression vectors, *pT3-EH1*α*H-myrAkt* (Addgene, 179909) was digested with NotI-HF (NEB, R3189S) and NcoI (NEB, R0193S) to generate a linearized backbone. The coding regions of *BMI1* and *Sall4* were amplified from *pT3-EH1*α*-BMI1* (Addgene, 31783) and *pENTR223*.*1-Sall4* (DNASU, MmCD00083348) using primer pairs listed in **Supplementary Table 3**. The amplified fragments were integrated into the linearized backbone using NEBuilder HiFi DNA Assembly Master Mix (NEB, E2621L), following the manufacturer’s instruction. Details of all constructs and primers used in this study are provided in **Supplementary Tables 3**.

### Cloning CRISPR sgRNA for the target genes

To generate the *SB-LSL-Cas9-sgBmi1* plasmid, we used the *SB-LSL-Cas9* plasmid, as described previously(29). The details of all constructs, primers, and sgRNA sequences used for this cloning are provided in **Supplementary Table 3**.

### Quantitative Reverse Transcriptase PCR (qRT-PCR)

Total RNA was extracted from frozen mouse liver tissues using RNeasy mini-Kit (Qiagen, 74106) and complementary DNA (cDNA) was synthesized using the PrimeScript™ RT Reagent Kit (TaKaRa, RR037A) according to the manufacturer’s instructions. qRT-PCR was performed using diluted cDNA, primer pairs (listed in **Supplementary Table 4**), and Power SYBR™ Green PCR Master Mix (ThermoFisher, 4367660) on a QuantStudioX real-time PCR system (Applied Biosystems). *Hprt* and *18S* were used as reference genes for normalization.

### Statistical Analysis

For all mouse experiments, sample size was pre-determined based on previous literature describing SB-HDTVI-mediated liver carcinogenesis(30). Littermates were randomized into experimental groups for HDTVI and maintained throughout the course of the experiment in a non-blinded manner. All subsequent molecular, immunohistochemical, and immunofluorescence analyses was conducted in a blinded manner. Variability in the data is presented as mean ± standard deviation (SD) in bar plots. Comparisons between two groups were performed using a two-tailed unpaired t-test. Statistical significance was defined as P<0.05. The significance thresholds were as follows: P<0.05 (^*^significant, ^*^), p<0.01 (^**^highly significant, ^**^), and p<0.005 (^***^extremely significant, ^***^), with additional asterisks denoting increasing levels of significance. All statistical analyses were performed using GraphPad Prism 10.0 (GraphPad Software).

### Data Availability Statement

The data generated in this study are available upon request from the corresponding author.

## RESULTS

### SALL4 is specifically required for *YAP1*-mediated HC-to-CCA transformation. (Fig. 1)

To investigate the role of SALL4 in liver cancer development, we first examined *Sall4* mRNA expression across diverse murine liver cancer models generated using SB-HDTVI approach. Given the validated *SALL4* expression in human CCA, subset of HCC and mixed and/or combined HCC-CCA, we have included following murine liver cancer subtypes in our assessment: murine CCA, HCC, combined HCC-CCA (cHCC-CCA), and mixed HCC/CCA (mHCC/CCA) generated by distinct oncogenic drivers (**Fig. 1A**). qPCR analysis revealed that *Sall4* expression is significantly elevated in *myrAkt-YAP1*^*S127A*^-driven CCA (*AY*-CCA), *myrAkt-NRAS*^*G12V*^-driven cHCC-CCA (*ANRAS*-cHCC-CCA), and *KRAS*^*G12D*^*-shp53*-driven mHCC/CCA (*KP53*-mHCC/CCA) compared to the healthy control livers. However, *Sall4* expression was undetectable in *myrAkt-N1ICD*-driven CCA (*AN*- CCA), *KRAS*^*G12D*^*-sg-p19*-driven CCA (*KP19*-CCA), *β-Cat*^*S45D*^*+Met*-HCC and *β-Cat*^*S45D*^*+Nrf2*^*G31A*^-HCC suggesting its context-dependent expression in murine liver cancer subtypes (**Fig. 1A**). To determine whether *Sall4* is required for tumor development in *AY*-CCA, *ANRAS-*cHCC-CCA, and *KP53-*mHCC/CCA models, we conditionally deleted *Sall4* in HCs–the cells in which oncogenes are hydrodynamically delivered to induce tumors. Specifically, we used *Sall4*^*(flox/flox)*^ mice and administered *adeno-associated virus serotype 8 (AAV8)-TBG-Cre* (1×10^12^ Genome Copies; GC, intravenously, i.v.) to generate HC-specific *Sall4* knockouts (*Sall4* KO^hepΔ^). Two weeks later, we initiated tumorigenesis by hydrodynamically delivering *AY, ANRAS, or KP53* to induce liver cancer (**Fig. 1B**). Mice that received *AAV8-TBG-GFP* were used as controls *(Sall4 WT*^*GFP*^). In the *ANRAS* and *KP53* models, loss of *Sall4* had no significant impact on tumor development. Both KO and WT mice exhibited comparable tumor burden, evident by gross observation, and comparable liver weight-to-body weight (LW/BW) ratios (**Fig. 1C, D**). Furthermore, immunohistochemistry (IHC) analysis for pan-cytokeratin (panCK, CC/CCA marker) and HNF4A (HC/HCC marker) revealed no significant change in the HCC/CCA composition between *Sall4* KO^hepΔ^ and *Sall4* WT^GFP^ mice (**Fig. 1E, F**). These findings suggest that *Sall4* is dispensable in *ANRAS-* or *KP53-*driven liver cancer development and tumor lineage commitment. In contrast, mice lacking *Sall4* in the *AY*-CCA exhibited a robust reduction in gross tumor burden, evidenced by a significantly lower LW/BW ratio and diminished macroscopic tumor growth compared to *Sall4* WT^GFP^ (**Fig. 1C, D**). Histologically, HA-tagged (myrAKT) tumor regions were drastically reduced in *Sall4* KO^hepΔ^ livers (**Fig. 1G**). Despite this reduction, the remaining tumors were still panCK^+^ but HNF4A^−^, indicating that *Sall4* deletion does not alter tumor cell fate but inhibits CCA tumor expansion (**Fig. 1E, G**). Collectively, these findings demonstrate that SALL4 is specifically required for *AY*-driven CCA formation but is dispensable for *ANRAS-* and *KP53-* driven tumorigenesis.

**Figure 1.**
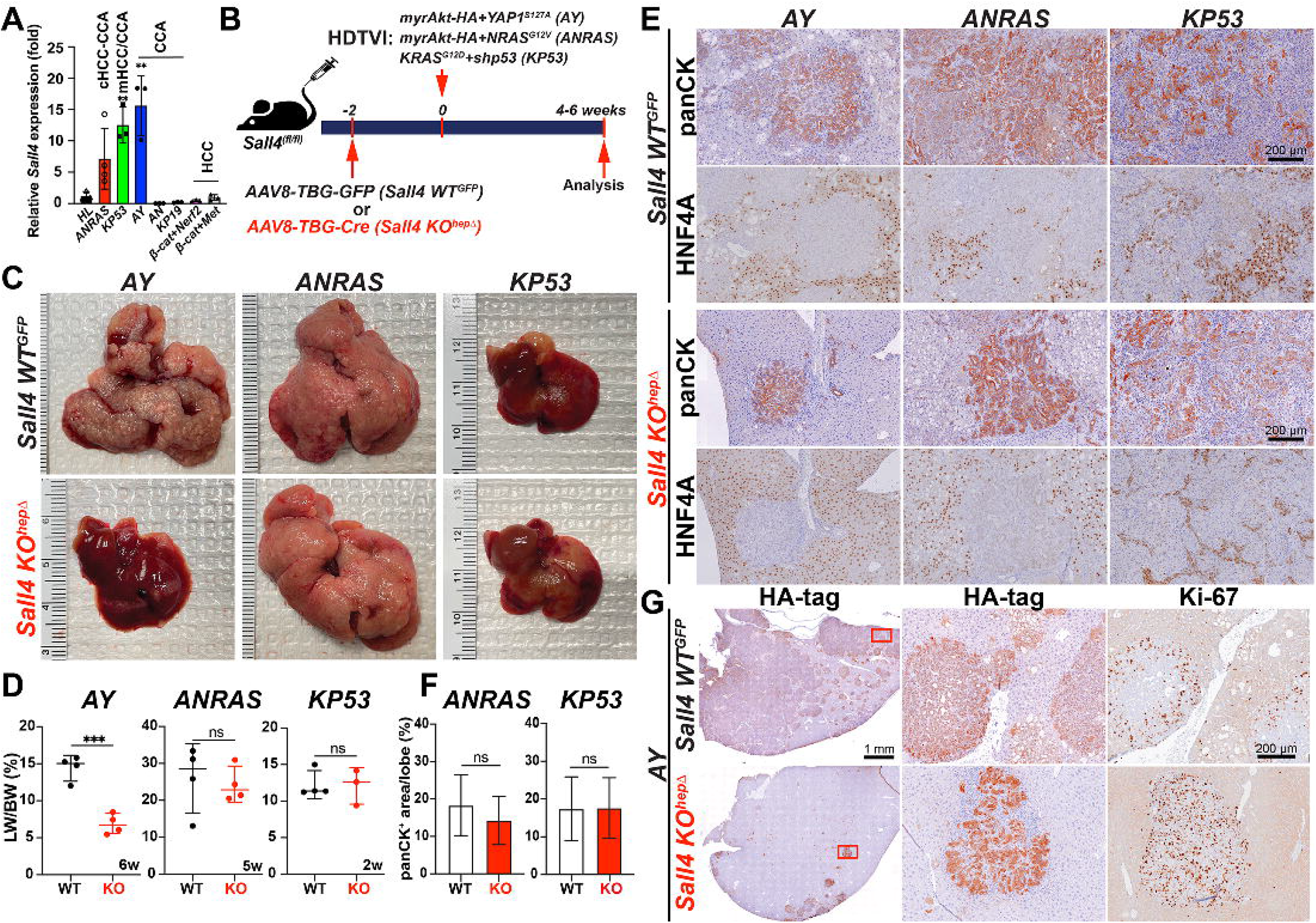
SALL4 is specifically required for *YAP1*-mediated HC-to-CCA transformation. (**A**) qRT-PCR analysis showing the relative expression levels of *Sall4* in different murine liver tumor models (*AY*-CCA, *ANRAS*-combined HCC-CCA, *KP53*-mixed HCC/CCA, and β*-catenin*-driven HCCs). (**B**) Schematic representation of the experimental design. HDTVI was used to deliver oncogenes (*AY*: *myrAkt-HA*+*YAP1*^*S127A*^; *ANRAS*: *myrAkt-HA*+*NRAS*^*G12V*^; *KP53*: *KRAS*^*G12D*^*+shp53*) into *Sall4*^*fl/fl*^ mice, combined with a prior intravenous injection (2 weeks before HDTVI) of *AAV8-TBG-GFP* (*Sall4 WT*^*GFP*^) or *AAV8-TBG-Cre* (*Sall4 KO*^*hep*Δ^). Livers were analyzed 4–6 weeks post-HDTVI. (**C**) Representative liver images from each model (*AY, ANRAS, KP53*) comparing *Sall4 WT*^*GFP*^ and *Sall4 KO*^*hep*Δ^. (**D**) LW/BW ratio analysis across all models comparing *Sall4 WT*^*GFP*^ and *Sall4 KO*^*hep*Δ^ (n = 3-4 mice per group). (**E**) IHC for panCK (cholangiocyte lineage marker) and HNF4A (hepatocyte lineage marker). (**F**) Quantification of panCK^+^ area (%) in *ANRAS* and *KP53* livers. (**G**) IHC for HA-tag and the proliferation marker Ki-67 in *AY* tumors from *Sall4 WT*^*GFP*^ and *Sall4 KO*^*hep*Δ^ mice. Data are presented as mean ± SD; Student’s t-test (^*^p < 0.05, ^**^p < 0.01, ^***^p < 0.001, ns = not significant). Scale bars: (E) 200 μm; (G) 1 mm (overview), 200 μm (magnified views).

### The loss of *Sall4* impairs HC-to-CCA transformation and its clonal expansion at the early stage. (Fig. 2)

To investigate the role of *Sall4* in the HC-to-CCA transformation, we explored histological changes in *Sall4* KO^hepΔ^ and *Sall4* WT^hep^ livers at the early stage in this model (at 2 weeks post-HDTVI) using IHC (**Fig. 2A**). Efficient deletion of *Sall4* in HCs was achieved using *AAV8-TBG-Cre;Sall4*^*(flox/flox)*^ strategy and confirmed by significant reduction in *Sall4* mRNA level in *Sall4* KO^hepΔ^ liver by qPCR assay (**Fig. 2A**). Notably *Sall4* KO^hepΔ^ livers exhibited a significant reduction in HA-tagged area suggesting reduced HC-to-CCA transformation and/or impaired viability of transfected cells (**Fig. 2B-C**). We first assessed proliferation and cell death of HA-tag^+^ cells using Ki-67 IHC and TUNEL staining. The Ki-67 proliferation index was significantly decreased in KO and TUNEL staining showed no notable difference in cell death between groups (**Fig. 2D-F**). These findings indicate that SALL4 is essential for the clonal expansion of *AY*-transfected cells but not required for their survival during the early phase of *AY*-driven CCA transformation. At this stage in WT livers, HA-tag^+^ *AY*- transfected HCs exhibited a transition toward LPC/CCA-like morphology, characterized by SOX9 and Ki-67 positivity, forming early-stage, clonally expanding nodular structures (**Fig. 2D, G**)(31). To further investigate HC-to-CCA transformation and the clonal expansion, we compared the number and size of SOX9^+^ CCA nodules between WT and KO livers, serving as indicator of HC-to-CCA conversion efficiency and subsequent proliferation respectively. (**Fig. 2G, H**). The deletion of *Sall4* led to a significant reduction in both parameters, suggesting that both HC-to-CCA reprogramming and their subsequent expansion was attenuated by *Sall4* deletion. Taken together, these findings suggest that SALL4 plays an essential role in both the initiation of HC-to-CCA reprogramming and following clonal expansion during the early tumorigenesis.

**Figure 2.**
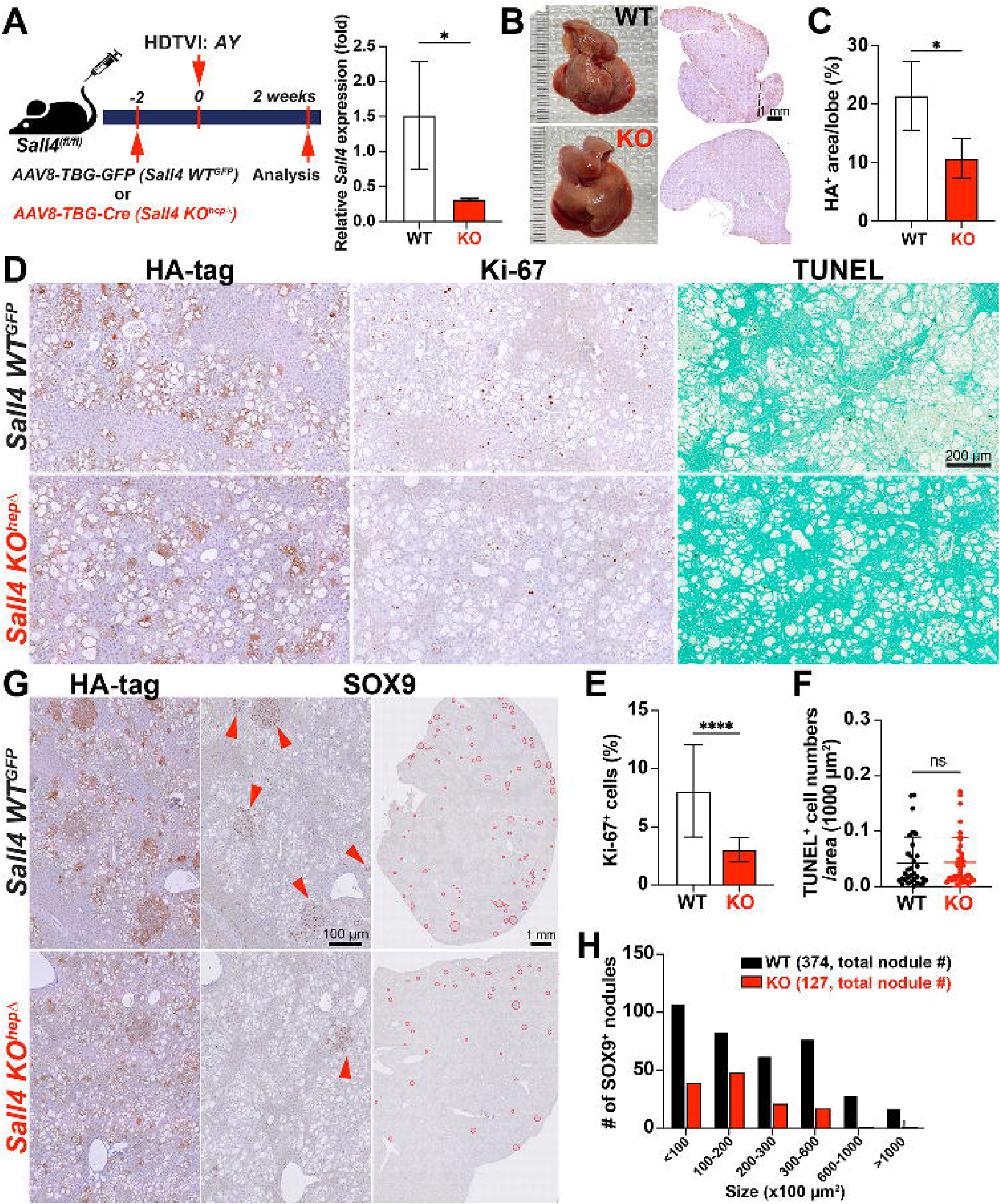
The loss of *Sall4* impairs HC-to-CCA transformation and its clonal expansion at the early stage. (**A**) Schematic representation of the experimental design. *Sall4*^*fl/fl*^ mice underwent HDTVI with transposon vectors expressing HA-tagged *myrAkt* and *YAP1*^*S127A*^ (*AY*). Mice received either *AAV8-TBG-GFP* (*Sall4 WT*^*GFP*^) or *AAV8-TBG-Cre* (*Sall4 KO*^*hep*Δ^) prior to HDTVI, leading to hepatocyte-specific *Sall4* knockout before tumor development. Livers were analyzed 2 weeks post-HDTVI. qRT-PCR analysis showed relative *Sall4* expression in *Sall4 WT*^*GFP*^ and *Sall4 KO*^*hep*Δ^ livers at 2 weeks post-HDTVI. (**B**) Representative images of livers and corresponding overview HA-tag IHC staining from *Sall4 WT*^*GFP*^ and *Sall4 KO*^*hep*Δ^ mice, showing differences in tumor burden. (**C**) Quantification of HA-tag^+^ area per lobe (%) in *Sall4 WT*^*GFP*^ and *Sall4 KO*^*hep*Δ^ mice. (**D**) IHC staining for HA-tag, Ki-67 and TUNEL staining. (**E-F**) Quantification of Ki-67^+^ proliferating cells (%) and TUNEL^+^ cells in *Sall4 WT*^*GFP*^ and *Sall4 KO*^*hep*Δ^ tumors. (**G**) IHC staining for HA-tag and SOX9. Red arrowheads highlight SOX9^+^ nodules in *Sall4 WT*^*GFP*^ tumors, reduced in *Sall4 KO*^*hep*Δ^ tumors. (**H**) Quantification of SOX9^+^ tumor nodules, categorized by size (x100 µm^2^), comparing *Sall4 WT*^*GFP*^ (black) and *Sall4 KO*^*hep*Δ^ (red) groups. The total number of nodules is indicated in parentheses. Data are presented as mean ± SD; Student’s t-test (^*^p < 0.05, ^****^p < 0.0001). Scale bars: (D) 200 µm; (G) 1 mm (overview), 100 µm (magnified views).

### *Sall4* overexpression disrupts *YAP1*-driven HC-to-CCA transformation while enhancing the clonal expansion of fatty LPC-like cells. (Fig. 3)

To further elucidate the role of *Sall4* in *AY*-CCA tumor formation, we investigated the effects of its gain-of-function *in vivo*. For this purpose, we conditionally overexpressed *Sall4* in *AY*-transfected HCs by co-delivering the *pT3-EF1*α*-Sall4-V5* plasmid with *AY* into FVB wild-type mice via HDTVI (hereafter referred to as *AY-Sall4*). Mice injected with *pT3-EF1*α*-GFP* (*AY-GFP*) served as controls. Animals were sacrificed at 2.5 and 5 weeks post-HDTVI to assess early reprogramming events and tumor progression, respectively (**Fig. 3A**). Successful *Sall4* overexpression was confirmed by qPCR at 2.5 weeks post-HDTVI (**Fig. 3B**). Notably, *AY*-Sall4 livers displayed prominent lipid accumulation, resulting in a significant increase in LW/BW ratio, which reached ∼8 compared to ∼6 in controls (**Fig. 3C-D**). At 2.5 weeks, IHC revealed striking histological differences between groups (**Fig. 3E**). In *AY*-GFP livers, early CCA nodules developed as HA-tag^+^;panCK^+^;Ki-67^+^;HNF4A^−^ clusters, consistent with early-stage reprogrammed CCA (**Fig. 3E-G**). In *AY*-GFP livers, approximately 20% of HA-tag^+^ cells expressed panCK but lacked HNF4A, indicating active HC-to-CCA transformation whereas remaining HA-tag^+^ cells in *AY*-GFP livers were panCK^−^;HNF4A^+^, displaying fatty HC morphology with Ki-67 positivity. In *AY*- *Sall4* livers, all HA-tag^+^ cells retained HNF4A expression and lacked panCK staining, indicating impaired early HC-to-CCA fate conversion (**Fig. 3E, G**). Proliferation of these HA-tag^+^;HNF4A^+^ cells were significantly decreased compared with HA-tag^+^ cell population in WT liver while accumulated excessive lipids, forming macro vesicular steatosis that displaced nuclei (**Fig. 3H, I**). This suggests expansion of a non-reprogrammed, lipid-rich LPC-like intermediate cell population. To further investigate steatosis, early liver injury and microenvironmental changes, we performed TUNEL staining and serum chemistry analysis at 2.5 weeks (**Fig. 3J–L**). TUNEL^+^ cell numbers were slightly but significantly increased in *AY*-*Sall4* livers, while there were no significant differences in serum aspartate aminotransferase (AST), alanine aminotransferase (ALT), alkaline phosphatase (ALP), or total bilirubin level, indicating limited functional hepato-biliary injury with cell death in *AY*-Sall4 livers. Interestingly, CD45^+^ immune cell infiltration and α-SMA^+^ myofibroblast activation showed significant decrease in *AY*-Sall4 livers compared to controls (**Fig. 3K, M-N**). These findings imply a diminished inflammatory and fibrotic microenvironment in *AY*-*Sall4* livers, despite ongoing cell death and steatosis. By 5 weeks post-HDTVI, *AY*-GFP livers exhibited grossly visible CCA tumors, with LW/BW ratios exceeding 10 (**Fig. 3C-D**). In contrast, *AY*-*Sall4* livers continued to show diffuse steatosis without tumor formation. The LW/BW ratio in *AY*-*Sall4* remained stable, similar to 2.5 weeks, and significantly lower than *AY*-GFP livers at 5 weeks. Histologically, panCK^+^; HNF4A^−^ CCA regions in *AY*-GFP livers expanded to occupy over 60% of the HA^+^ area by 5 weeks, whereas in *AY*-*Sall4*, only ∼20% of HA-tag^+^ area expressed weak panCK and still co-expressed HNF4A, suggesting an incomplete reprogramming LPC-like intermediate cell status (**Fig. 3G-I, O**). Although hepatic fat content was slightly reduced in *AY-Sall4* livers by this time point, fully reprogrammed HNF4A^−^ CCA nodules remained absent. Given that transient AKT-driven steatosis typically resolves as HCs undergo CCA fate conversion(31, 32), these findings indicate that *Sall4* overexpression interferes with HC-to-CCA fate switching by sustaining a progenitor-like, lipid-rich state. This intermediate state is characterized by co-expression of HNF4A and panCK, high proliferation, and impaired terminal differentiation. Collectively, these results demonstrate that *Sall4* overexpression inhibits CCA progression in the *AY* model by multiple mechanisms including impairing HC-to-CCA lineage conversion, promoting clonal expansion of proliferative, steatotic, LPC-like intermediate cells, and enhancing death of *AY*-transfected cells. These findings underscore the distinct and opposing roles of SALL4 gain- and loss-of-function in cellular reprogramming and YAP1- mediated cholangiocarcinogenesis.

**Figure 3.**
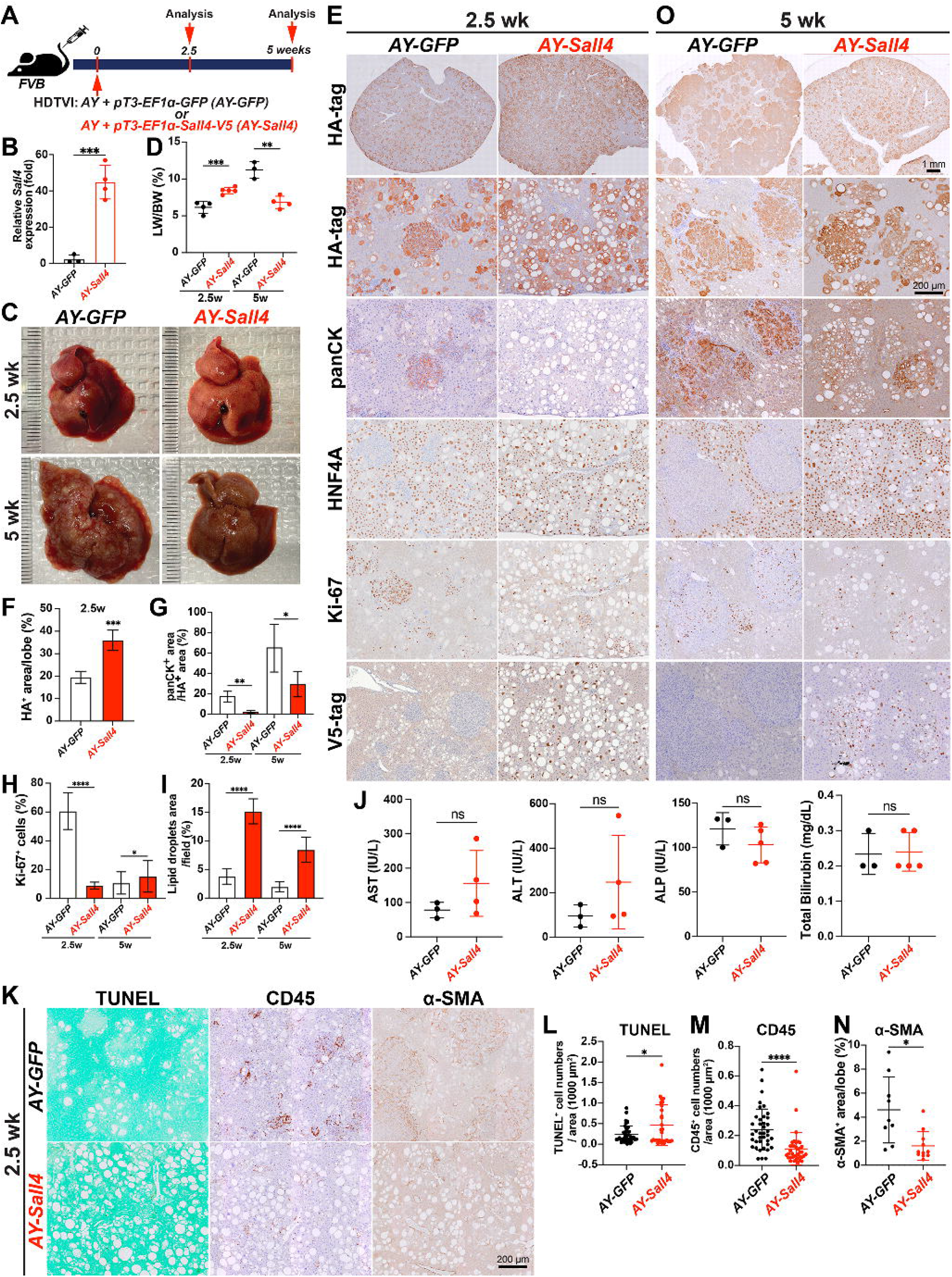
*Sall4* overexpression disrupts *YAP1*-driven HC-to-CCA transformation while enhancing the clonal expansion of fatty LPC-like cells. (**A**) Schematic representation of the experimental design. FVB mice underwent HDTVI with transposon vectors expressing HA-tagged *myrAkt* and *Yap1*^*S127A*^ (*AY*) combined with GFP expression (*AY-GFP*) or V5-tagged *Sall4* (*AY-Sall4*) plasmids and were analyzed at indicated time points (2.5 weeks and 5 weeks post-HDTVI). (**B**) qRT-PCR showing *Sall4* expression in *AY-Sall4* compared to *AY-GFP*. (**C**) Representative liver images from *AY-GFP* or *AY-Sall4* mice at 2.5 (top) and 5 weeks (bottom) post-HDTVI. (**D**) LW/BW ratio analysis comparing *AY-GFP* and *AY-Sall4* at indicated time points. (**E, O**) IHC analysis for indicated markers (HA-tag, panCK, HNF4A, Ki-67, V5-tag) at at indicated time points. (**F-N**) Quantification of indicated parameters (tumor area, panCK^+^ area, lipid droplet area, Ki-67^+^ cells, AST, ALT, ALP, bilirubin, CD45, TUNEL, α-SMA) comparing *AY- GFP* and *AY-Sall4*. Data are presented as mean ± SD; Student’s t-test (^*^p < 0.05, ^**^p < 0.01, ^***^p < 0.001, ^****^p < 0.0001). Scale bars: (E, O) 1 mm (overview), 200 μm (high magnification); (K) 200 μm.

### *Bmi1* modulation phenocopied *Sall4* modulation in *AY*-driven HC-to-CCA tumorigenesis. (Fig. 4)

Given that BMI1 is a validated functional SALL4 downstream effector in diverse malignancies including liver cancer(24), we first examined whether *Bmi1* expression is elevated in *AY-Sall4* liver compared with *AY-GFP* control livers. qPCR analysis showed a significant upregulation of *Bmi1* mRNA expression by *Sall4* overexpression in *AY-Sall4* livers (**Fig. 4A**), supporting further functional analysis of *Bmi1* modulation in the *AY*-CCA model. Along with *Sall4*, we explored the effects of *Bmi1* loss- and gain-of-function in the *AY*-CCA model. To conditionally eliminate *Bmi1* in the *AY*-CCA model, we used the previously validated *SB-Cas9- sgRNA* system(29). We delivered *SB-Cas9-sgBmi1* plasmid, along with *AY*, into FVB wild-type mice via HDTVI (*Bmi1 KO*^*CCA*Δ^), and evaluated tumor development at 6-8 weeks post-injection. As controls, we used *SB-Cas9- sgEmpty* plasmid under the same experimental conditions (*Bmi1 WT*^*CCA*^) (**Fig. 4B**). Similar to *AY-Sall4 KO*^*hep*Δ^, tumor-specific *Bmi1* ablation significantly impaired *AY*-CCA formation, as assessed by prominently reduced gross CCA formation and LW/BW ratio (**Fig. 4C-D**). In the control group, lethal CCA tumors (HA- tag^+^;panCK^+^;HNF4A^−^) exhibited LW/BW ratios reaching 20. In contrast, very few but highly proliferating CCA nodules developed in the *Bmi1 KO*^*CCA*Δ^, with LW/BW ratios around 5, comparable to healthy controls (**Fig. 4C-F**). This observation indicate that *Bmi1* is indispensable for *AY*-CCA development similar with *Sall4*. Next, to examine the effect of *Bmi1* overexpression in *AY*-CCA, we injected *pT3-EF1*□*-GFP or pT3-EF1*□*-BMI1-V5* plasmids along with *AY* plasmids and evaluated tumorigenesis at 6-8 weeks post-HDTVI (**Fig. 4B**) and successful *Bmi1* upregulation was confirmed by qPCR (**Fig. 4G**). Interestingly, LW/BW ratios and gross observation showed that *Bmi1* overexpression robustly reduces CCA tumor burden (**Fig. 4D, H**). IHC analysis revealed that HA-tag^+^ cells were foamy HCs with lipid accumulation, with only a few panCK^+^;HNF4A^−^ CCA tumor nodules (**Fig. 4I, J**). These findings were consistent with the earlier comparison of *Sall4* overexpression and control in *AY*-CCA at 6 weeks (**Fig. 3D-E**). Along with the elevated transcription levels of *Bmi1* in *Sall4*- overexpressing livers (**Fig. 4A**), these data suggests that *Sall4* overexpression may disrupt HC-to-CCA transformation through BMI1, leading to a phenocopy of *Bmi1* overexpression. In summary, the similar effects of gain and loss of function for both *Sall4* and *Bmi1* in *AY*-CCA suggest that BMI1 may act as a downstream effector of SALL4 in *AY*-CCA tumor development.

**Figure 4.**
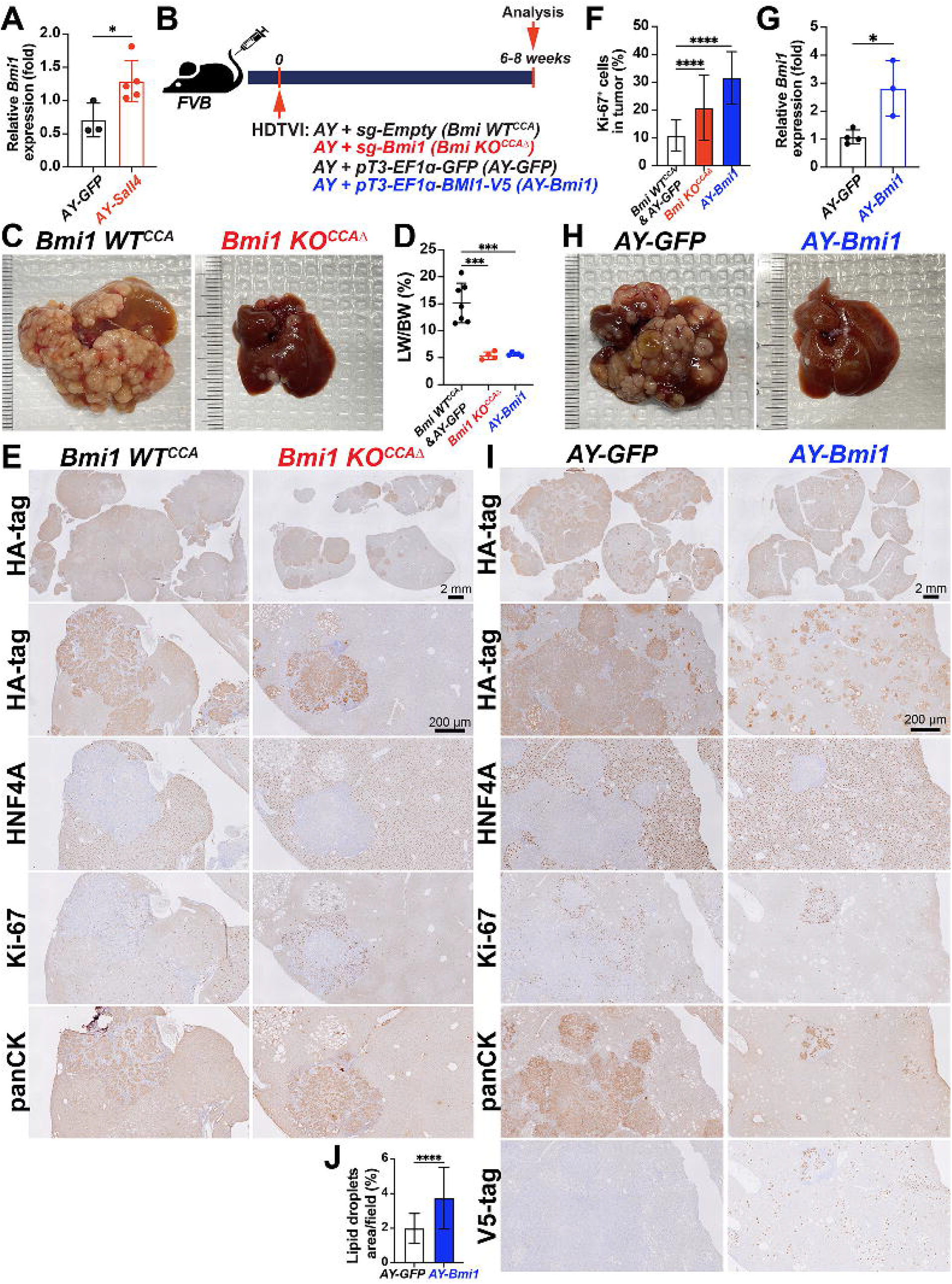
*Bmi1* modulation phenocopied *Sall4* modulation in *AY*-driven HC-to-CCA tumorigenesis. (**A**) qRT-PCR analysis showing *Bmi1* expresson in *AY-Sall4* compared to *AY-GFP*. (**B**) Schematic experimental design. (**C, H**) Representative macroscopic images of livers harvested from indicated groups. (**D**) LW/BW ratio comparison (n = 3–5 per group). (**E, I**) IHC analysis for indicated markers (HA-tag, HNF4A, Ki-67, panCK, V5-tag) in experimental groups. (**F, J**) Quantification of Ki-67^+^ proliferating cells (%) and lipid droplets area per field (%) comparing *Bmi1 KO*^*CCA*Δ^ and *AY-Bmi1* tumors. (G) qRT-PCR analysis showing *Bmi1* expression in *AY-Bmi1* compared to *AY-GFP*. Data are presented as mean ± SD; Student’s t-test (^*^p < 0.05, ^***^p < 0.001, ^****^p < 0.0001). Scale bars: (E, I) 2 mm (overview), 200 µm (magnified views).

### SALL4 regulates regenerative HC-to-CC conversion in the DDC-fed cholestasis model. (Fig. 5)

Given the critical role of SALL4 in *YAP1*-dependent HC-to-CCA malignant reprogramming, we next explored its function in non-malignant reparative HC-to-CC reprogramming using the validated DDC-fed murine cholestasis model with regenerative HC-to-CC conversion(5, 33). To enable transient HC lineage tracing alongside HC-specific *Sall4* deletion, we administered *AAV8-TBG-GFP* and *AAV8-TBG-Cre* virus into *Sall4*^*(f/f)*^ mice (i.v., 1×10^12^ GC) followed by 4 weeks of DDC feeding (*Sall4 KO*^*hep*Δ*GFP*^). As a control, *Sall4*^*(f/f)*^ mice receiving only *AAV8-TBG-GFP* were used to allow HC lineage tracing while maintaining intact *Sall4* expression (*Sall4 WT*^*hep-GFP*^) (**Fig. 5A**). Serum biochemistry analysis at 4 weeks of DDC diet revealed no significant differences in ALP, ALT, and AST levels between *Sall4 WT*^*hep-GFP*^ and *Sall4 KO*^*hep*Δ*GFP*^ mice (**Fig. 5B**). However, total bilirubin levels were significantly lower in the *Sall4 KO*^*hep*Δ*GFP*^ group, suggesting that while HC damage was comparable, *Sall4* deletion in HCs may help mitigate bile duct injury caused by DDC diet feeding. To further assess the impact of HC-specific *Sall4* loss on the hepatic microenvironment in the DDC model, we evaluated CD45^+^ immune cell infiltration and fibrosis by IHC. Interestingly, both immune infiltration and fibrotic response were significantly increased in *Sall4*-deleted livers (**Fig. 5C–E**), despite the decreased biliary injury as indicated by serum total bilirubin (**Fig. 5B)**. These counterintuitive findings suggest that *Sall4* in HCs may influence the hepatic microenvironment through non-cell-autonomous manner, possibly via paracrine signaling in the setting of cholestatic injury. To evaluate HC-to-CC transdifferentiation in the absence of *Sall4*, we investigate both early phase HC-to-LPC activation (ductular reaction) and subsequent LPC-to-CC differentiation respectively through histological analyses. The ductular reaction, characterized by bile duct proliferation and LPC activation(34), was assessed via SOX9 (marking both CC and LPC) and Ki-67 IHC. Despite reduced total bilirubin levels, *Sall4 KO*^*hep*Δ*GFP*^ livers exhibited a significantly enhanced ductular reaction, as evidenced by significantly increased number of SOX9^+^ cells and Ki-67^+^ proliferating cells (**Fig. 5F–H**). Since the ductular reaction in this model involves expansion of pre-existing CC, CC-derived LPCs, and HC-derived LPCs(35), we next specifically assessed the HC-derived LPC population. Co-immunofluorescence (IF) for GFP (HC lineage tracing marker), HNF4A (mature HC marker), and SOX9 (LPC/CC marker) revealed a significant increase in SOX9^+^;HNF4A^+^;GFP^+^ HC-derived LPCs in *Sall4 KO*^*hep*Δ*GFP*^ livers compared to controls, suggesting that *Sall4* loss in HCs may promote the initial transition of HCs into LPCs under cholestatic conditions (**Fig. 5K, L**). To determine whether HC-derived LPCs in *Sall4 KO*^*hep*Δ*GFP*^ livers could fully differentiate into mature CCs, we assessed GFP^+^ HC-derived CC conversion using co-IF for CK19, a mature CC marker. Consistent with robust increase in initial ductular reaction and HC-to-LPC formation in *Sall4 KO*^*hep*Δ*GFP*^ livers (**Fig. 5F-H**), the total number of CK19^+^ mature CCs was higher in *Sall4 KO*^*hep*Δ*GFP*^ livers compared to controls (**Fig. 5M-N**). However, the proportion of GFP^+^;CK19^+^ HC-derived cells among a total of CK19^+^mature CCs was significantly reduced in *Sall4 KO*^*hep*Δ*GFP*^ livers (**Fig. 5M, O**), implying that enhanced CK19^+^ CCs in portal area is likely due to the expansion of pre-existing CCs rather than HC-to-CC transdifferentiation. These findings indicate that *Sall4* loss enhances initial HC-to-LPC activation but impairs their differentiation into mature, functional CCs in cholestatic livers. This imbalance likely contributes to an expansion of pre-existing CCs, which may compensate for bile duct injury and explain the reduced total bilirubin levels observed in *Sall4 KO*^*hep*Δ*GFP*^ mice fed DDC diet.

**Figure 5.**
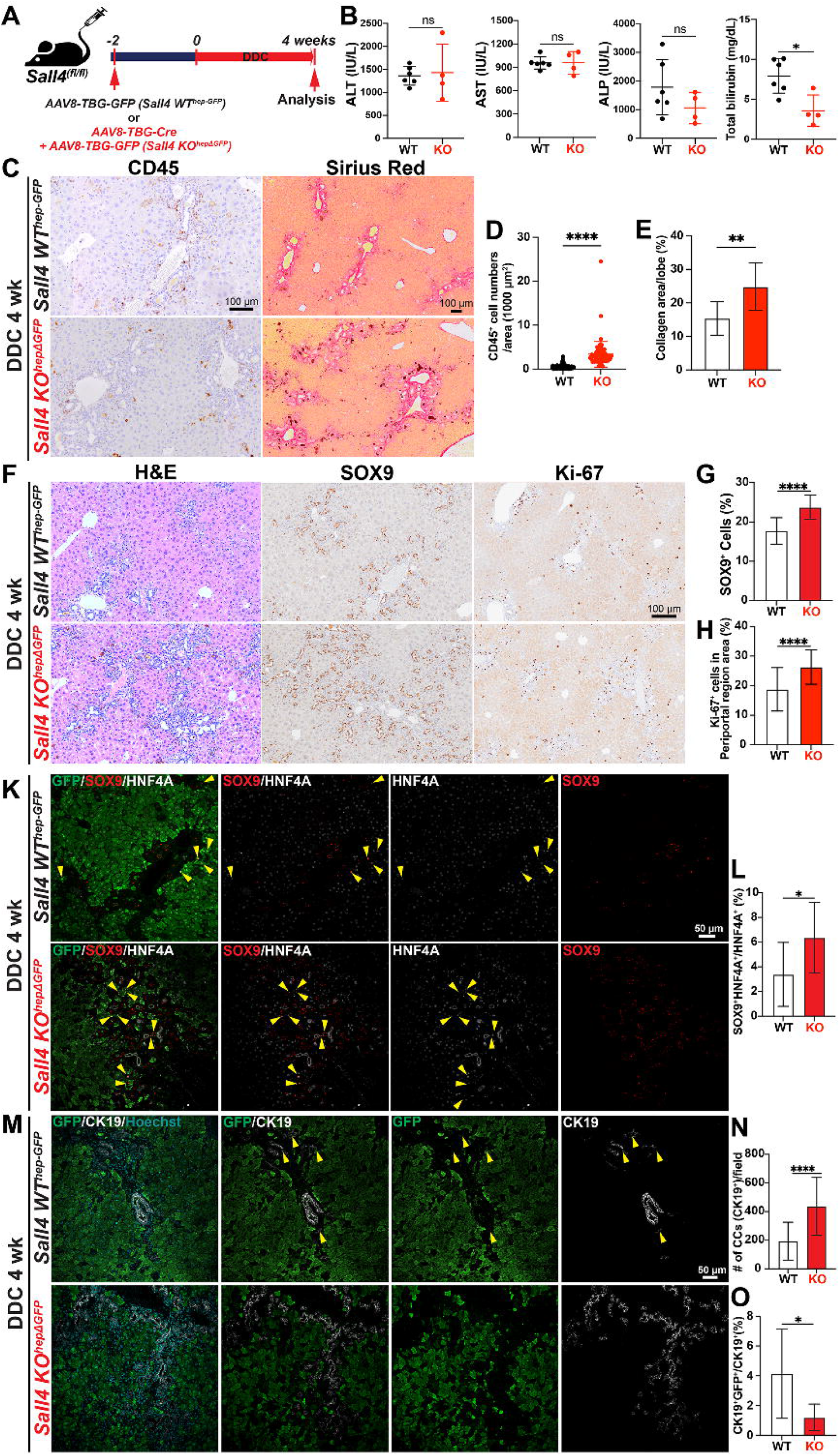
SALL4 regulates regenerative HC-to-CC conversion in the DDC-fed cholestasis model. (**A**) Schematic experimental design. *Sall4*^*fl/fl*^ mice were injected with either *AAV8-TBG-GFP* (*Sall4 WT*^*hep-GFP*^) or *AAV8-TBG-Cre* (*Sall4 KO*^*hep*Δ*GFP*^) 2 weeks prior administration of a DDC-containing diet for 4 weeks. Liver tissues were analyzed at the end of the treatment period. (**B**) LW/BW ratio comparison and serum biochemistry analysis comparing ALT, AST, ALP, and total bilirubin levels between *Sall4 WT*^*hep-GFP*^ and *Sall4 KO*^*hep*Δ*GFP*^ groups (n = 4–6 mice per group). (**C**) Representative images of CD45 IHC and Sirius Red staining in livers sections from both groups. (**D-E**) Quantification of (D) CD45^+^ inflammatory cells per 1000 µm^2^ and (E) collagen deposition (%) by Sirius Red staining. (**F**) Representative liver sections stained with H&E, SOX9 (cholangiocyte lineage marker), and Ki-67 (proliferation marker). (**G-H**) Quantification of (G) SOX9^+^ cells (%) and (H) Ki-67^+^ cells in periportal area (%). (**K**) IF staining for SOX9, HNF4A, and GFP. Yellow arrowheads indicate intermediate cells expressing both cholangiocyte (SOX9) and hepatocyte (HNF4A) markers. (**L**) Quantification of SOX9^+^HNF4A^+^ double-positive imtermediate cells among total HNF4A^+^ cells (%). (**M**) IF staining for CK19 and GFP to visualize hepatocyte-derived cholangiocyte. (**N**) Quantification of CK19^+^ cholangiocytes per microscopic field. (**O**) Quantification of CK19^+^GFP^+^ double-positive cells among total CK19^+^ cells (%). Data are presented as mean ± SD; Student’s t-test (^*^p < 0.05, ^**^p < 0.01 ^****^p < 0.0001, ns = not significant). Scale bars: (C-F) 100 µm; (K-M) 50 µm.

## DISCUSSION

Our study identifies SALL4 as a crucial regulator in YAP1-mediated HC-to-CCA malignant transformation and regenerative HC-to-CC conversion in cholestasis model. Furthermore, our data suggest that BMI1, validated downstream effector of SALL4, mimics its role in the malignant transformation, reinforcing a mechanistic link between YAP1, SALL4 and BMI1 in liver tumorigenesis (**Fig. 6**).

**Figure 6.**
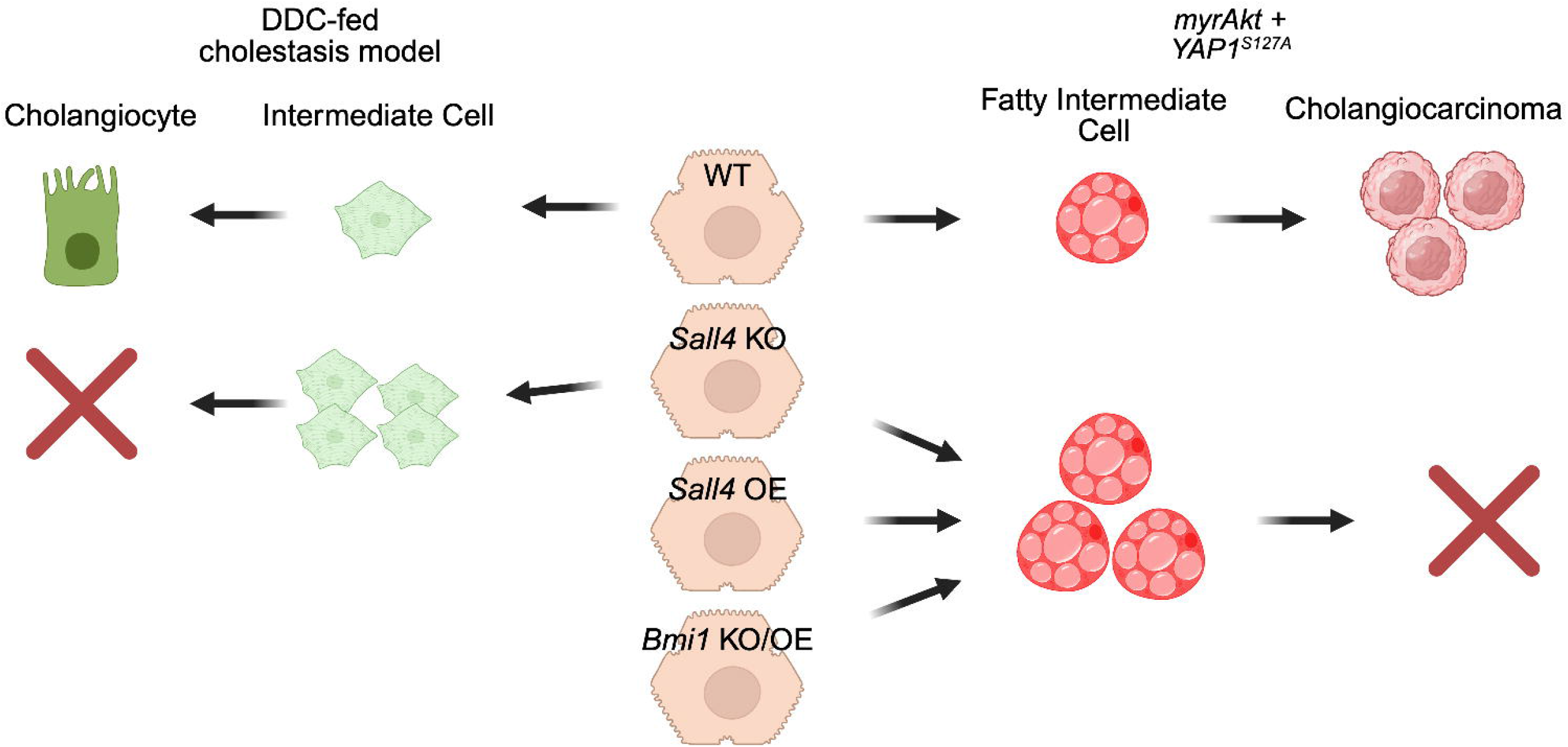
Graphical summary illustrating the role of SALL4 and BMI1 in reparative HC-to-CC conversion in DDC-induced cholestatic liver injury and *AY*-driven malignant HC-to-CCA transformation.

Interestingly, the function of SALL4 appears to be specific to YAP1-driven CCA tumorigenesis, as it is dispensable in other liver cancer models driven by *KRAS, NRAS*, or *p53*, regardless of tumor lineage fate. This suggests that the role of SALL4 is context-dependent and may be restricted to tumors where YAP1 signaling is dominant, irrespective of histologic classification. Consistently, another key finding is that SALL4 is neither detectable nor functional in the *AN*-CCA model, despite YAP1 being required for transformation in this setting(10), further supporting the idea that SALL4 engagement within the YAP1 pathway depends on additional oncogenic or epigenetic contexts. This specificity may indicate that while YAP1 activation is necessary for HC-to-CCA conversion(10), SALL4 is selectively recruited in particular developmental or tumorigenic programs. Clinically, SALL4 expression is enriched in combined or mixed HCC-CCA tumors, which display both hepatocytic and CC features reminiscent of bipotential progenitor-like cells such as hepatoblasts and LPCs(18). Given the established roles of YAP1 in bile duct development and HC dedifferentiation, these findings suggest that SALL4 operates primarily within the YAP1 signaling axis in both malignant and regenerative contexts. Supporting this, prior studies have shown co-expression of YAP1 and SALL4 in gastric cancer stem cells, suggesting a cooperative role in lineage commitment during tumorigenesis(36). Therefore, the clinical relevance of SALL4 in cHCC-CCA likely reflects a subset of tumors characterized by YAP1 pathway activation, emphasizing the molecular heterogeneity of cHCC-CCA and the need to further dissect the YAP1– SALL4–BMI1 axis and its integration with other signaling networks in liver cancer.

Our gain-of-function experiments revealed an unexpected inhibitory effect of *Sall4* overexpression on HC-to-CCA transformation. Rather than promoting tumor formation, *Sall4* overexpression led to an expansion of an intermediate LPC-like cell population, characterized by persistent lipid accumulation(31) and dual HNF4A- panCK expression. This suggests that fine-tuned *Sall4* expression regulation in a delicate balance is critical for *AY-*driven HC dedifferentiation during AY-CCA transformation. In other words, excessive *Sall4* expression may stall LPCs in an intermediate state, thereby preventing their maturation into fully transformed CCA cells.

Interestingly, the AKT-mediated lipid accumulation observed in *Sall4*-overexpressing intermediate cells is likely due to impaired differentiation into the biliary lineage, rather than a direct effect of SALL4 on lipid metabolism. This paradoxical effect is reminiscent of biphasic role of SALL4 in stem cell biology, where it is required for early lineage commitment but must be tightly regulated to prevent aberrant differentiation.

Although *BMI1* is highly expressed in HCC(37), and overexpression of *BMI1* together with *NRAS*^*G12V*^ can induce well-differentiated HCC tumors in mice(37), its role in lineage conversion and CCA tumorigenesis remains unclear. Notably, our findings show that SALL4 and BMI1 share similar phenotypic effects–loss of either impairs CCA formation, while overexpression promotes the expansion of LPC-like populations. These observations suggest that BMI1 mediates at least part of function of SALL4 in YAP1-driven HC-to-CCA transformation. Furthermore, the fact that *Sall4* overexpression leads to *Bmi1* upregulation indicates a direct transcriptional relationship between these two factors. Given that BMI1 is a core component of the Polycomb Repressive Complex 1 (PRC1)(38, 39), it is possible that SALL4-BMI1 signaling modulates epigenetic programs critical for lineage conversion. Future studies should investigate whether SALL4 directly regulates *Bmi1* transcription or acts through chromatin remodeling mechanisms.

Beyond its role in tumorigenesis, we examined the role of SALL4 in regenerative HC reprogramming using a DDC-fed cholestasis model. Notably, pan-HC-specific *Sall4* ablation resulted in a unique response to cholestatic injury; this robustly stimulated the ductular reaction while reducing bile duct injury. This contradicts previous assumptions that ductular reaction positively correlates with liver disease severity(40-42), suggesting that SALL4 in HC may regulate ductal proliferation via a paracrine mechanism during the injury. In these perspectives, to clarify the cell type-specific roles of SALL4, it would be interesting to investigate cell-type specific knockout models, such as *OPN-Cre*^*ERT2*^ (biliary-specific) or *Alb-Cre* (simultaneous HC and CC deletion) under cholestatic conditions. This could help determine how SALL4 functions in HCs and/or CCs in liver disease.

Furthermore, unlike malignant transformation, where *Sall4* loss impairs HC-to-CCA oncogenic reprogramming, its loss in cholestatic injury promotes HC transition into LPCs (co-expressing HNF4A and SOX9). This is similar to the effects of *Sall4* overexpression in the *AY*-CCA model, although AKT-mediated lipid accumulation is absent. These findings indicate that SALL4 plays distinct roles in malignant versus injury-induced HC reprogramming, operating through different molecular mechanisms. These results suggest that *Sall4* deletion might facilitate the reverse process: CC/LPC-to-HC transdifferentiation, opposite to HC-to-CC conversion. This possibility could be investigated using dual-lineage tracing system (simultaneously tracking HC and CC lineages) in a mouse model, beyond the debating of leakiness on murine CC lineage tracing system.

Given the established involvement of TRAF3/NIK(43), Notch(11), and TGF-β(44) signaling pathways in HC-to-CC fate conversion, as well as prior evidence supporting a SALL4–TGF-β signaling axis, we investigated whether SALL4 modulates these pathways. Using qPCR, we assessed the expression of key target genes in healthy liver, *AY*-CCA, and *AY-Sall4-*CCA liver tissues (**Fig. S1**). Interestingly, no significant changes were observed in the expression of *Traf3, NIK*, or the Notch pathway targets *Hey1* and *Hes1*. Among TGF-β targets, only *Timp3* was significantly downregulated in *AY-Sall4-*CCA livers, whereas *Serpine1* levels remained unchanged. These findings suggest that SALL4 may suppress TGF-β signaling in the *AY*-CCA context. Although previous studies have implicated a functional SALL4–TGF-β axis(45, 46), the observed reduction in TGF-β target expression may instead reflect decreased fibrotic activation associated with reduced CCA burden, rather than direct transcriptional crosstalk.

Given its established role in cholestasis and liver cancer, SALL4 may provide therapeutic potential while its context-dependent effects necessitate a precise approach with caution such as complete inhibition of SALL4 could prevent malignant transformation but may also impair normal liver regeneration. The success of implication may largely rely on the development of fine-tune delivery strategy rather than systemic regulation of SALL4 entirely could offer a more effective strategy. Additionally, the identification of BMI1 as a downstream effector provides an alternative therapeutic avenue. BMI1 inhibitors, either alone or in combination with YAP1 inhibitors, may effectively block HC-to-CCA transformation. Given the growing clinical interest in YAP1- targeted therapies, understanding how SALL4 and BMI1 interact within the YAP1 signaling pathway may help refine treatment strategies for aggressive liver cancers.

In conclusion, our study highlights SALL4 as a pivotal regulator of HC reprogramming, with distinct roles in promoting early transformation while restraining full CCA progression in a context-dependent manner. The identification of BMI1 as a downstream effector further elucidates the mechanistic function of SALL4 in YAP1- driven tumorigenesis. These findings enhance our insights into the molecular mechanism of HC plasticity and may offer future investigations into potential therapeutic strategies targeting this pathway in liver cancer subtype and other liver diseases.

## ACKNOWLEDGEMENT

Work performed in the Pitt Biospecimen Core (RRID:SCR_025229) and services and instruments used in this project were supported, in part, by the University of Pittsburgh, the Office of the Senior Vice Chancellor for Health Sciences. We also thank Drs. Satdarshan (Paul) Monga and Brandon Lehrich for kind sharing cDNA derived from murine HCC samples.

---

**Supplementary Figure 1. SALL4 overexpression suppresses *Timp3* expression but has limited impact on Notch- and TRAF3/NIK–related genes in AY-driven CCA.**

qRT-PCR analysis for indicated genes related to the Notch (*Hes1, Hey1*), TRAF3/NIK (*Traf3, Map3k14*) and TGF-β (*SerpinE1, Timp3*) pathways in liver tissues from healthy liver (HL), *AY-GFP*, and *AY-Sall4* groups at 2.5 wk post-HDTVI. Relative mRNA expression levels were normalized to housekeeping genes (*Gapdh*) and presented as fold changes relative to HL. SALL4 overexpression significantly reduced *Timp3* expression compared to the AY-GFP group (^*^p < 0.05). No significant differences (ns) were observed in the expression of the other genes. Data are shown as mean ± CD (n = 3–5 per group). Statistical significance was determined by Student’s t-test.

